# Molecular principles of Piezo1 activation by increased membrane tension

**DOI:** 10.1101/823518

**Authors:** Dario De Vecchis, David J Beech, Antreas C Kalli

## Abstract

Piezo1 is a mechanosensitive channel involved in many cellular functions and responsible for sensing shear-stress and pressure forces in cells1–3. Piezo1 plays a critical role in the circulatory system and tissue development. Mutations on Piezo1 are linked to human diseases such as lymphedema2,4 or hematological disorders such as hemolytic anaemia5 and resistance to malaria6. Hypotheses for Piezo1 gating include the “force-from-lipids” principle7,8 that suggests that Piezo1 senses mechanical forces through the bilayer1,9 and a direct involvement of the cytoskeleton as well as the extracellular matrix in Piezo1 activation10,11. However, the molecular and structural changes underpinning the Piezo1 gating mechanism and how the channel senses forces in the membrane remain unknown. Here we reveal the activation mechanism of Piezo1 and the structural rearrangements that occur when Piezo1 moves from a closed to an open state when mechanical tension is applied to the cell membrane. Our results show that Piezo1’s curved shape is stable in a native-like model membrane without tension creating a membrane indentation with a trilobed topology. Upon stretching Piezo1 adapts to the stretched bilayer by flattening and expansion of its blade region. In our simulations Piezo1 expands up to a planar circular area of ∼680 nm^*2*^ comparable with previous structural data and hypotheses12–14. Piezo1 flattening and expansion results in changes in the beam helix tilt angle. These movements result in the tilting and lateral movement of the pore lining TM37 and TM38 helices. This leads to the opening of the channel and to the movement of lipids that occupy Piezo1 pore region outside of this region, revealing for the first time the structural changes that happen during Piezo1 mechanical activation. The changes in the blade region are transmitted to helices TM37 and 38 via hydrophobic interactions and by interactions of neighbouring subunits via the elbow region. The flat structure of Piezo1 identified in this study exposes the C-terminal extracellular domain (CED) that in the closed state is hidden in the membrane and presumably from shear stress. Our results provide new structural data for different states of Piezo1 and suggest the molecular principles by which mechanical force opens the Piezo1 channel, thus coupling force to physiological effect via ion permeation.

---

To investigate how mechanical forces act on the Piezo1 channel, we simulated the recent Piezo1 structure solved by cryo-electron microscopy (cryo-EM) in a model membrane using a serial multi-scale molecular dynamics (MD) simulation approach. Missing residues were added to the structure prior to the simulations (see Methods). An initial coarse-grained (CG) simulation of 500 ns was carried out to equilibrate the bilayer around Piezo1 and create the unique Piezo1 footprint suggested previously12–14. Although the CG approach was critical for creating the Piezo1/membrane system, it does not allow us to study the Piezo1 dynamics. For this reason, the system was backmapped to an atomistic resolution and further simulated in independent repeat simulations applying pressure to the bilayer (i.e. along x and y axes) ranging from the native-like pressure of +1 bar to -40 bar as described in Methods. Under these conditions, we were able to calculate the lateral pressure (*P*_*L*_) applied on the bilayer which is 14.2 mN/m for the -5 bar, 45.8 mN/m for the -20 bar, 59 mN/m for the -30 bar and 67.8 ± 0.35 mN/m for the -40 bar system. The asymmetric bilayer used in our simulations mimics the native endothelial membrane in which Piezo1 functions15 (see Methods).

## Tension causes Piezo1 to flatten and expand

The different pressures cause expansion of the simulated box to a different degree with the membrane in the system adapting to each pressure. This results in an increase of the area per lipid (APL) in both leaflets for all systems; the APL for the system with the highest tension in the bilayer was 90.09 ± 0.84 Å^2^ and 89.46 ± 0.96 Å^2^ the upper and the lower leaflets, respectively (see Supplemental Table S1 for all APLs). A similar approach revealed that an increase of the APL resulted in the transition of the TREK-2 mechanosensitive channel for a down to an up conformation16.

Our analysis above shows that the APL reaches a plateau for both leaflets within the first 10 ns of the simulation. Therefore, each system was simulated for at least 50 ns to allow sufficient time for Piezo1 to equilibrate in the bilayer. Indeed, our analysis shows that the opening of the channel occurs after the first 10 ns of simulation in conjunction with the increase of the APL (Fig. 4a, b and Supplemental Table S1). However, to further understand if more extended simulation time alters the APL, two of our systems the -30 bar and the +1 bar systems were extended to 100 ns. In both cases the analysis confirms than no change in the APL is observed in the longer simulations (Supplemental Table S1). Moreover, a further repeat simulation at -40 bar was carried out obtaining a comparable result, thus indicating that the systems are equilibrated, and the simulated time is sufficient to study the Piezo1 structural transitions under tension. The Piezo1 structure shows a remarkable adaptation to a stretched bilayer with its N-terminal blades, which remain embedded within the bilayer during all simulations, to extend towards more flatten conformations (Fig. 1a). The degree of flattening was different for the different tensions with the system with the highest tension resulting in an almost flat Piezo1 structure (Fig. 1a). Fig. 1b shows the correlation between the area of the simulated box and the root mean-square deviation (RMSD) calculated using the Cα atoms of the blades, demonstrating that they both increase until the bilayer stops expanding. Note that in the bilayer without tension (+1 bar) Piezo1 maintains its curved shape even though this simulation is twice as long (i.e. 100 ns) as some of the simulations with a tension applied to the bilayer. This shows that the curved shape of Piezo1 blades forces the bilayer to adopt a curved trilobed shape. Interestingly, whilst Piezo1 reaches a flat conformation in both the -30 bar and -40 bar systems, larger conformational changes with more opening of the pore are observed in the -40 bar simulations. Even when the -30 bar systems were extended up to 100 ns (compared to 50 ns of the -40), the structural changes in this system were smaller (Fig. 1a, b), suggesting that there is a cross-talk between Piezo1 and bilayer with Piezo1 sensing forces from the bilayer after it reaches the flat conformation. Our RMSD analysis shows the Piezo1 blades as the major contributors to the RMSD drift, whereas the CED mostly retained its initial conformation with comparable values between the systems (Fig. 1c).

**Figure 1.**
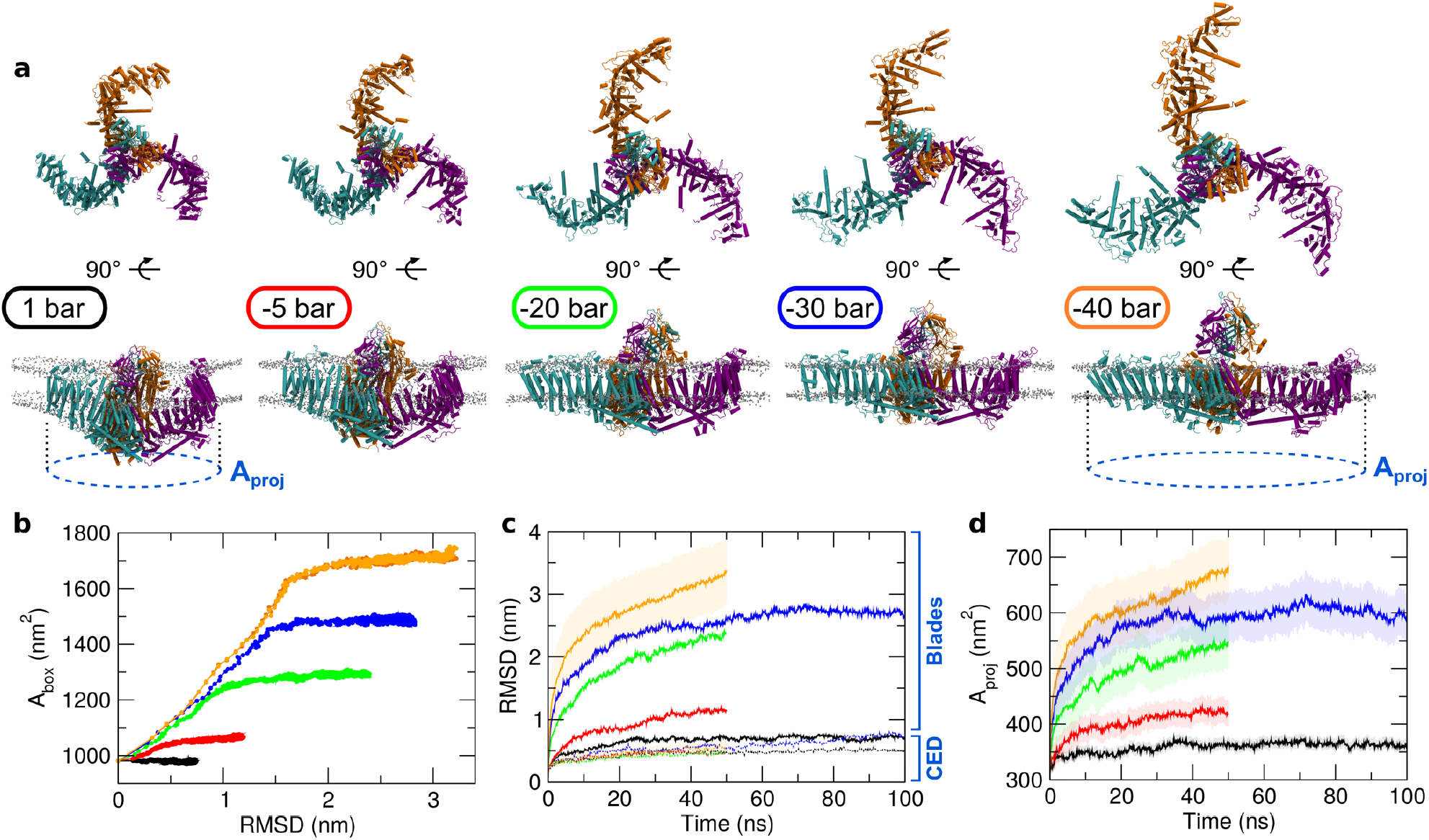
Piezo1 channel flattens and its CED becomes exposed in response to increased membrane tension. For each system, the applied negative pressure in each simulation is indicated with different colours: black, 1 bar; red, -5 bar; green, -20 bar; blue, -30 bar; orange, -40 bar. (**a**) Snapshots of the last frame from simulations of Piezo1 embedded in a model membrane bilayer. (**b**) Correlation between the area, i.e. x-side × y-side, of the simulated box (A_box_) and the root mean-square deviation (RMSD) calculated using the protein Cα of the Piezo1 blades. (**c**) RMSD calculated using the protein Cα of the C-terminal extracellular domain (CED), dashed lines (lower values), and of the rest of the protein (i.e. the Piezo1 blades), straight lines (higher values). (**d**) Projected area of the Piezo1 membrane indentation on the membrane plane (A_proj_, indicated by a dashed blue circle in **a**) as a function of time. For the -40 system the timeline is the average between the two repeats. The standard deviation is indicated.

In addition, our results suggest that upon flattening Piezo1 also expands thus increasing the in-plane projection of the Piezo1 membrane indentation (A_proj_; Fig 1a). This is in good agreement with recent studies that also suggested an increase of the A_proj_ involved in the Piezo1, and recently Piezo2, activation^12–14^. The A_proj_ can be approximated by a circle. By approximating the Piezo1 triskelion to a regular triangle, we calculated the side of this triangle to further determine the circumscribed circle (i.e. the A_proj_) as described in the Methods. Our approximation is justified by the moderate standard deviation values of the side of the triskelion (i.e. the side of the triangle) which is ± 0.23 nm for the +1 bar, ± 0.53 nm for the -5 bar, ± 0.95 nm for the -20 bar, ± 0.88 nm for the -30 bar and ± 1.07 nm and ± 1.23 nm for the two repeats at -40 bar. This result is also indicative of a concurrent movement of the Piezo1 blades that flatten at the same time. Fig. 1d shows how the increase of applied mechanical force during simulations (e.g. more tension), causes an increase of the A_proj_ values which reach ∼680 nm^2^ at -40 bar (Fig. 1d, orange line). Recent cryo-EM structural data of the full-length Piezo2, estimated a A_proj_ of the dome to 700 nm^2^ 14. Although the current value for the full-length Piezo1 is still unknown, this agreement shows that upon flattening Piezo1 covers the area that is within the dome in the closed state14. This is in good agreement with the suggestion that the dome provides an energy store for Piezo1 activation12. We also note that in our atomistic simulations, even the +1 bar system shows a mild increase of the A_proj_ with respect to the initial coordinates from the cryo-EM (Fig. 1d, black line). Overall, our approach provides novel molecular insights into the activation and dynamics of Piezo1 in a native-like lipid environment in which this channel functions. Moreover, we dynamically quantify the different A_proj_ expansions under different mechanical stress and the area covered by a flat activated Piezo1 channel. Finally, the extended structure obtained at - 40 bar is corroborated by recent structural data which positively correlate the size of the vesicle with a flattened Piezo1 structure13. Flattening of Piezo1 may also be needed for Yoda1 binding^17^.

## Structural rearrangements in Piezo1 pore

Piezo1 flattening caused by membrane tension results in the displacement of the CED relative to the membrane plane (Fig 2a). Although our simulations did not show any changes in the position of the CED domain relative to the transmembrane (TM) domain, CED is hidden within the dome at the beginning of the simulations or in the system without tension applied in the bilayer. Flattening of Piezo1 and of the bilayer results in the exposure of CED as it emerges outside the dome. The degree of CED exposure above the dome increases as the tension in the bilayer increases; in the -30 bar and -40 bar systems in which a flat Piezo1 was reached CED is fully exposed. The density analysis also revealed that a considerable portion of the Piezo1 structure remained consistently exposed to the cytoplasm even after the expansion (Fig 2a). It is possible that these cytoplasmic regions remain exposed during activation providing a platform for Piezo1 interactions with the cytoskeleton, which is in agreement with its mechanoprotective role in Piezo1 function9.

**Figure 2.**
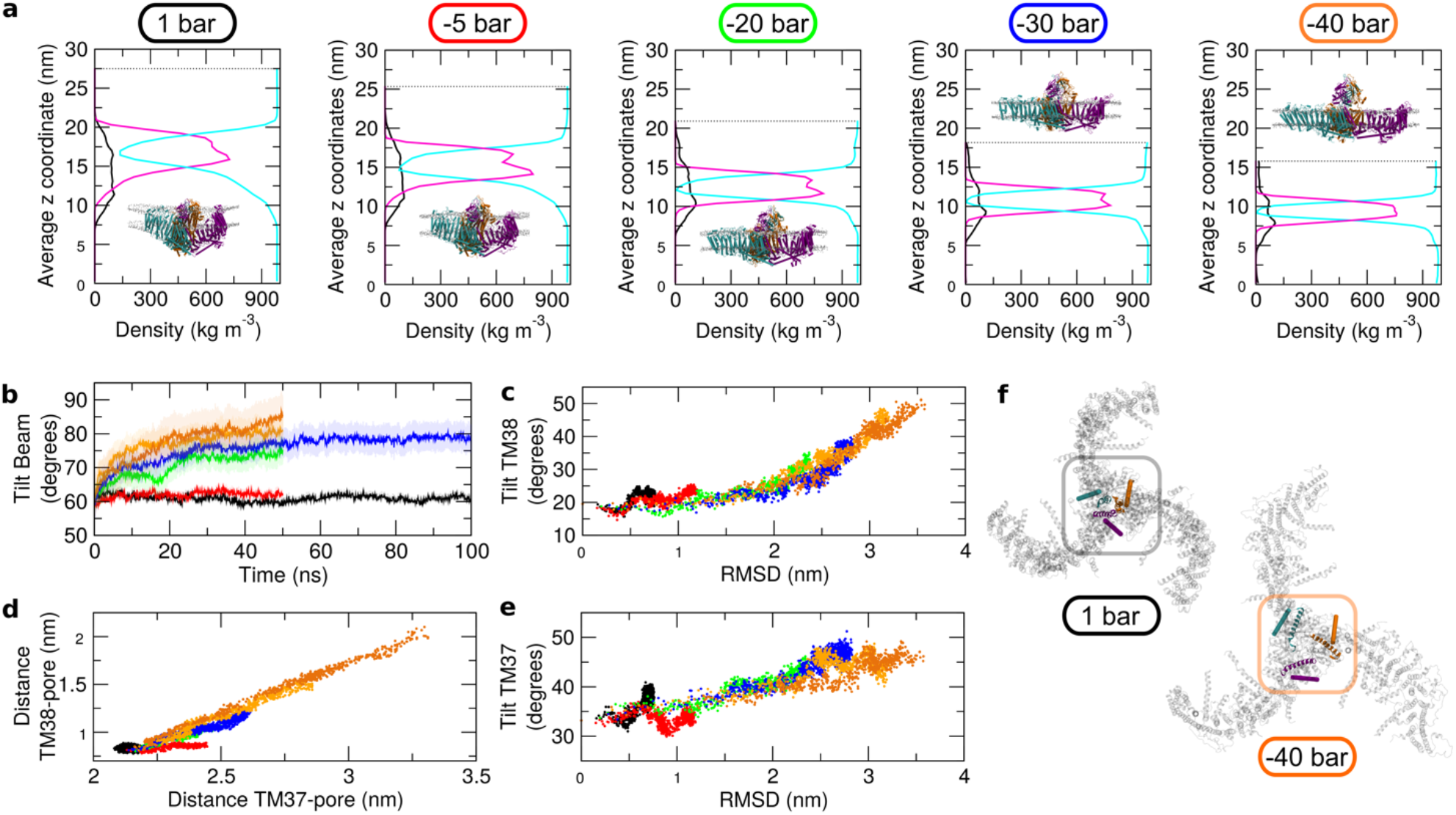
Coordinated tilting of Piezo1 channel beams and ion pore helices in response to increased membrane tension. (**a**) Partial density profile along the z direction (that is perpendicular to the bilayer x,y plane) for the protein (black line), the membrane (magenta line) and the solvent (i.e. water and ions, cyan line). Values from the first repeat at -40 bar are shown here. Insets are the last frames from simulations as shown in Fig. 1a. (**b**) Tilt angle relative to the bilayer normal for the beam helix. (**c**) Correlation between the RMSD of the Piezo1 blades and the tilt angle relative to the bilayer normal of the TM38 helix. (**d**) Correlation between the distances from the centre of mass of the TM38 or TM37 helices and the Piezo1 pore (**e**) Correlation between the RMSD of the Piezo1 blades and the tilt angle relative to the bilayer normal of the TM37 helix (**f**) Snapshots of the last frame from the +1 bar and -40 bar systems with the Piezo1 triskelion shown from the extracellular side. The centre of the triskelion is highlighted. The TM38 (cartoon) and the TM37 (ribbon) from each Piezo1 chain are also shown. Membrane and solvent are not shown for clarity. The colour code for the applied negative pressure in each simulation is indicated and is the same as in Fig. 1.

Another Piezo1 component which undergoes noticeable structural rearrangement is the beam helix (residues 1300-1365; Fig. 2b). Our data reveal that during the simulations with tension in the bilayer there is a change in the tilting angle of each Piezo1 beam (going from ∼65° to ∼85°), until reaching a position that is almost parallel to the bilayer surface when Piezo1 is in a flat conformation. Interestingly, the beam tilt movement is correlated with the change in the RMSD (Supplemental Fig. S1a) suggesting that this movement contributes to Piezo1 flattening. The beam is a long α-helix exposed to the cytoplasm which underpasses a great portion of each blade, precisely the three C-terminal proximal bundles. Therefore, our data suggest that each Piezo1 beam may act as a set of long levers and their concerted movement during mechanical sensing may possibly contribute to gating.

The precise molecular details of Piezo1 gating are still largely unknown. The protein is also characterised by a fast inactivation phase after gating^18,19^. Recent observations proposed a functional gate possibly regulating Piezo1 inactivation^20^. This gate is composed by the two conserved hydrophobic residues Leu2475 and Val2476 in the pore-lining TM helix 38^20^. Calculation of the tilt angle of the TM38 relative to the bilayer normal shows that higher negative pressure, and thus larger tension, corresponds to a higher tilting of the pore-lining TM38 (Supplemental Fig. S1b). The highest tilt angle is observed in the -40 bar system (Fig. 2c). Calculation of the distance of TM38 relative to the pore centre of mass also shows that larger tension, corresponds to a larger displacement (Supplemental Fig. S1c). Therefore, during flattening and expansion TM38 both increases its tilt angle and its distance from the pore centre of mass. The change in the TM38 tilting and distance from the channel pore is correlated with the changes in the RMSD calculated for the Piezo1 blades (Fig. 2c and Supplemental Fig. S1c). This shows that TM38 displacement/tilting occurs whilst Piezo1 flattens and expands (Fig. 2f). Interestingly, we found that the distance from the pore and tilting of the TM38 also correlates with the distance from the pore and tilting of the TM37 (Fig. 2d, e and Supplemental Fig. S1d, e), which is located anti-parallel (Fig. 2f). In conclusion, our simulations suggest a simultaneous lateral movement and tilting of the TM38 and TM37 that results in the opening of the pore.

## Tension leads to Piezo1 channel opening

Calculation of the water-filled cavity in Piezo1 pore region shows that the volume of this cavity increases with the applied negative pressures in our simulations until it reaches ∼75000 Å^3^ in the -40 bar (Fig. 3a). In our calculation the cavity is defined by the three TM38 (one from each subunit) and also some regions of the CED (see Methods), because the loops that connect this domain with the TM37/38 form a structure similar to a “cage” just above the channel mouth which is characterised by a funnel-shape (Fig. 3c). Fig. 3a also shows that tension of -30 bar on the membrane, although it contributes to flatting of Piezo1 structure and almost maximise the beam orientation, is not sufficient to open the channel pore and maximise the volume of the cavity; the cavity in the -30 system is not hydrated as much as in the -40 bar (Fig. 3b, d). Note that in the simulations of the system with no tension to the bilayer (1 bar) the cavity did not increased and the channel was not hydrated. We also found that lipid and cholesterol molecules initially overlap with this volume and therefore they are within the Piezo1 pore, but they increasingly leave the cavity as higher tension is applied to the bilayer (Fig. 4a). Three independent system set-ups followed by three independent CG equilibrations demonstrate that lipids get within the channel mouth during the equilibration phase. This demonstrates that there is enough space in the current closed structure of Piezo1 for lipids to occupy the channel mouth. This may also suggest that the presence of lipids in the pore may be required in order to maintain the channel mouth closed. Our results show that a higher tension to the membrane (that increases the APL in the proximity of the channel mouth), results in higher volume in the pore cavity as it enables lipids and cholesterol that occupy the channel mouth to move outside of it in addition to a larger movement of the pore lining helices. This hypothesis may explain biochemical data which suggest a functional role of lipids in Piezo1 regulation^21–23^. Therefore, our simulations suggest that the tension applied to the bilayer during functional activation, results not only in the displacement of TM38 and TM37 but also in the movement of lipids outside the Piezo1 pore. These two events occur simultaneously and may both be required for Piezo1 full opening.

**Figure 3.**
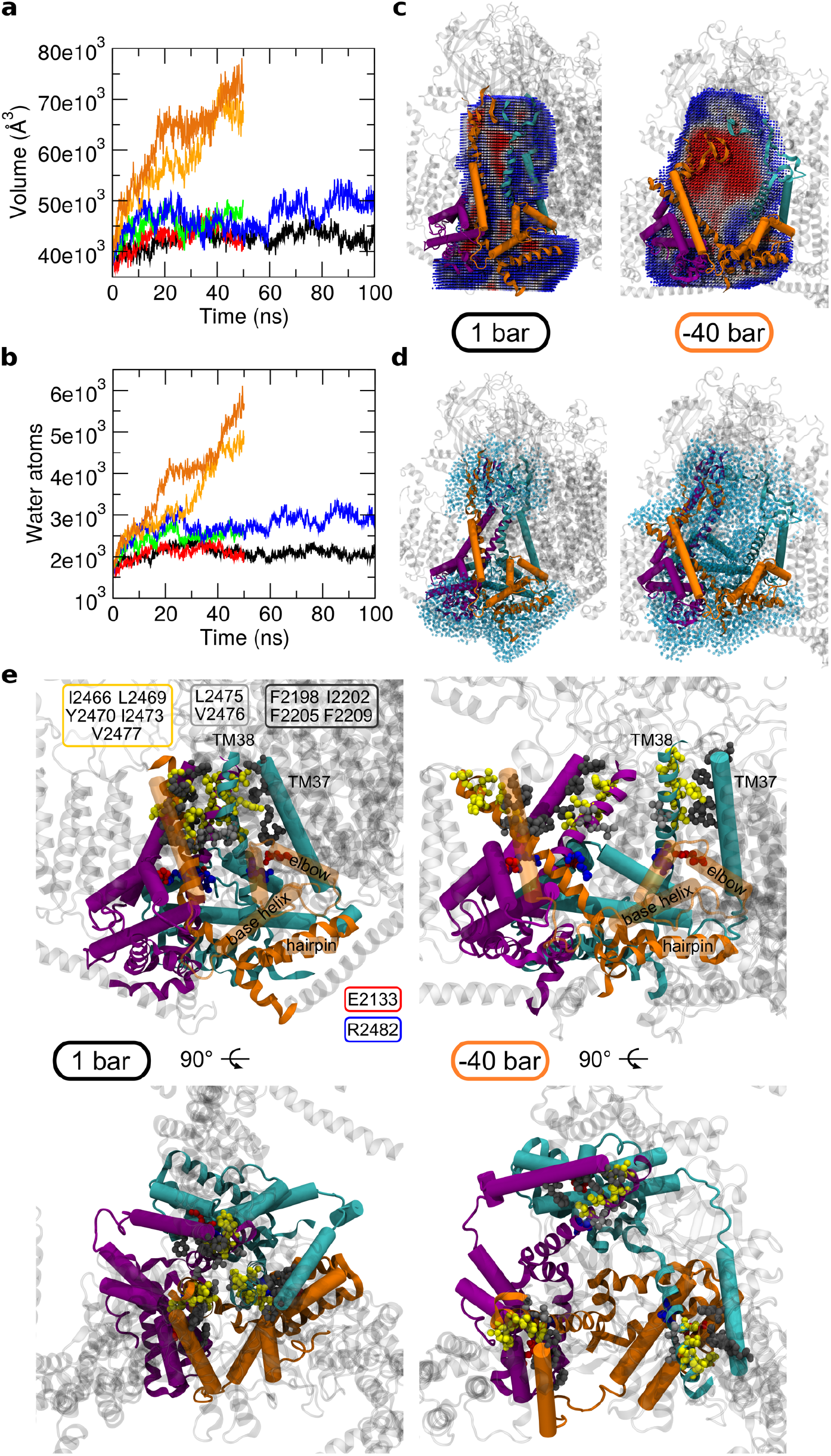
Expansion of the Piezo1 channel pore in response to increased membrane tension. (**a**) Volume of the cavity of the Piezo1 pore region as a function of time for each simulated system. The applied negative pressure in each simulation is indicated with different colours: black, 1 bar; red, -5 bar; green, -20 bar; blue, -30 bar; orange, -40 bar. (**b**) Number of water molecules overlapping with the calculated cavity in **a** as a function of time for each simulated system. (**c**) Volume of the cavity within the core of the Piezo1 structure, for the 1 bar and the -40 bar systems. The structures are the last frames from each simulation and show the Piezo1 pore region. For the -40 bar system the second repeat simulation is shown. The cavity (sliced) is shown as spheres and coloured according to the occupancy during the trajectory, from less populated (blue) to more populated (red). Residues selected for the calculation of the cavity are shown in ribbon (including the TM38 helix, see Methods). TM37, elbow and base helix are represented as cartoon cylinders. The Piezo1 chains are shown in the same colours as in Fig. 1a. (**d**) Water molecules within 8 Å from the residues selected for the calculation of the cavity for the 1 bar and the -40 bar systems. Water molecules are depicted in light cyan transparent spheres. The structures are the last frames from simulations and show the Piezo1 pore region. For the -40 bar system the second repeat is shown. The protein orientation and colour code are the same as in **c**. (**e**) Detail of the Piezo1 channel mouth for the 1 bar and the -40 systems shown from a side (above) and a extracellular (below) view. The structures are the same shown in **c** and **d** and show the hydrophobic interactions between the pore-lining helices TM38 (yellow residues) and the TM37 (dark grey residues). The salt bridge between the elbow and the TM38 from a neighbouring subunit is also shown (red and blue residues). Residues from the Piezo1 hydrophobic gate (residues L2475, V2476) are in light grey.

**Figure 4.**
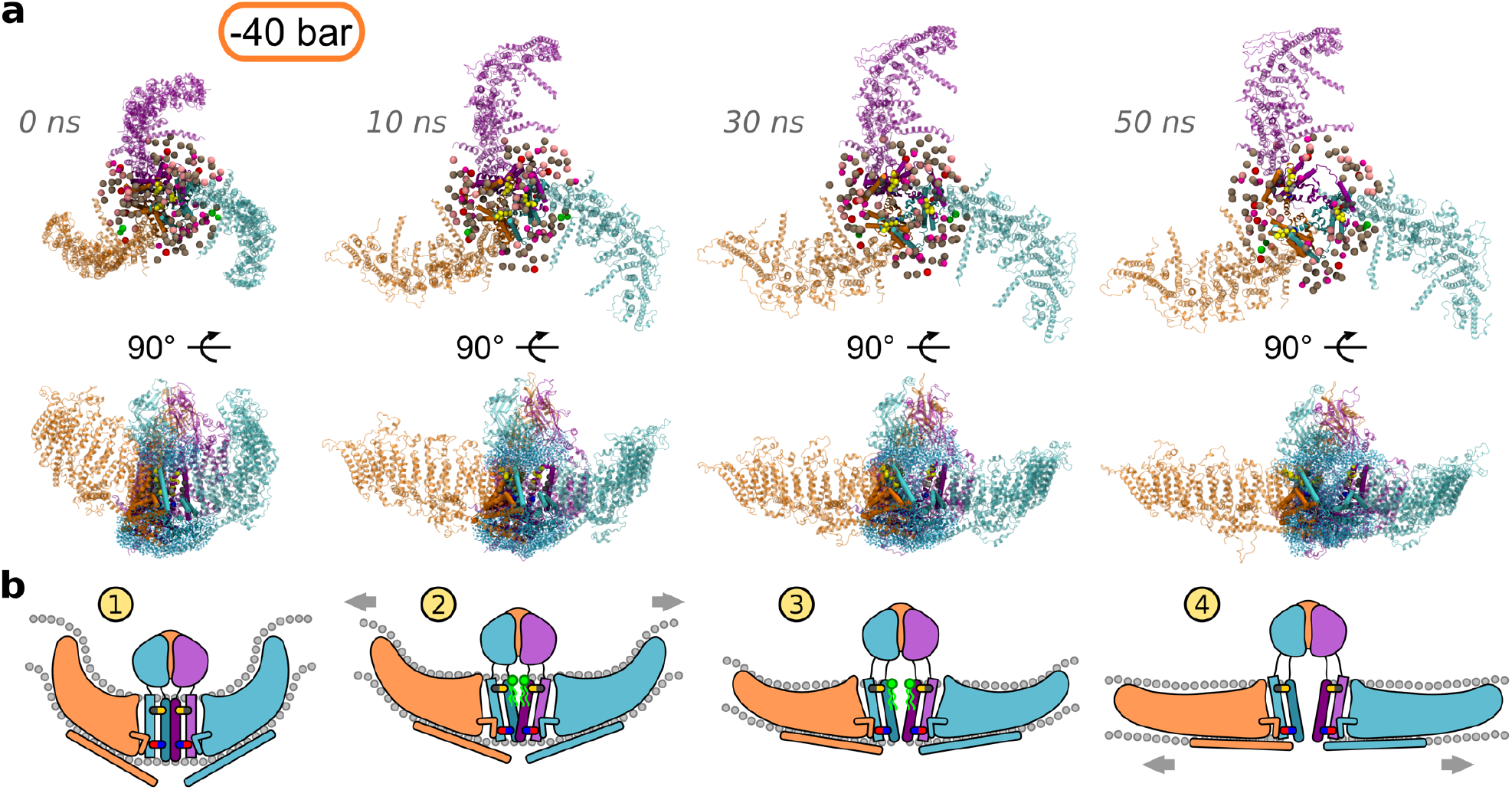
Proposed mechanism for the Piezo1 channel opening in response to membrane tension. (**a**) Snapshots from the second repeat simulation at -40 bar with the associated simulation time are shown. Membrane bilayer and solvent (i.e. water and ions) are not shown for clarity. The three Piezo1 subunits are indicated with different colours as in Fig. 1a. In the extracellular view at the top, the CED domain has been removed for clarity. Phosphorous and oxigen atoms from lipid headgroups and cholesterol respectively are shown (only lipids that are within 20 Å from the residues selected for the calculation of the Piezo1 pore region cavity; see Methods). The colour code is: POPC, tan; POPE, pink; POPS, red; PIP_2_, green; cholesterol, magenta. The Cα atoms from interacting residues between the helices TM37, TM38 and the elbow are indicated as spheres as in Fig. 3e. In the Piezo1 structures in the middle panel, water molecules within 8 Å from the residues selected for the calculation of the cavity (see Methods) are shown as light cyan transparent spheres. (**b**) Cartoon representation explaining the Piezo1 activation mechanism coupled to membrane tension. The colour code is the same as in **a**. The hydrophobic interactions between the TM37 and TM38 and the salt bridge between TM38 and the elbow from a neighbouring subunit, are indicated in yellow/grey and in blue/red, respectively. Lipids that initially occupy the Piezo1 channel mouth and progressively move away from it with the increase tension, are schematically depicted in green. Membrane tension and Piezo1 blade expansion are indicated by grey arrows.

An open question is how the mechanical stimuli are transferred from the blades to the channel pore. Our results suggest that the flattening and expansion of the blades is transmitted to the TM helices 37 and 38 guiding the pore opening. Noticeably, the pore region of the homologous Piezo2, has a remarkably similar structure14 which lead us to postulate a similar mechanism. In Piezo1 the pore-lining TM38, is located parallel, although with an opposite orientation, to the TM37. TM37 together with the elbow region (residues 2116-2142), a base helix (residues 2149 to 2175) and hairpin helices (residues 2501 to 2534) from all three subunits, create a cuff that encloses the pore-lining TM3812,13 (Fig. 3e). Noticeably, in all our simulations we noticed two main pivotal interactions: the first is the interaction between the TM37 and TM38 via the hydrophobic residues Ile2466, Leu2469, Tyr2470, Ile2473 and Val2477, that are adjacent to the aforementioned hydrophobic functional gate in TM38 (residues Leu2475 and Val2476^20^) and the residue Phe2198, Ile2202, Phe2205 and Phe2209 from the TM37 (Fig. 3e). Therefore, extension and expansion of the blades cause the displacement of TM37 that in turn “pulls” the pore-lining helix TM38 due their hydrophobic interactions. This results in the opening of the channel and might explain why in the -40 bar simulation, in which more expansion was observed, the pore was more open (Fig. 3e).

The second pivotal interactions occur in a region towards the C-terminal end of TM37 and TM38. Previous work suggested the importance of the negative charge at position Glu2133 (Glu2416 in Piezo2) for conductance and ion selectivity^24^. In particular, the single mutant E2133A showed half the unitary conductance with respect to the wildtype phenotype and the negative charge from the glutamate has been suggested to allosterically modulate the selectivity of the Piezo1 filter^24^. The Piezo1 cryo-EM structure allowed to precisely map the position of the Glu2133, which is located in a helix named elbow. Interestingly, the tip of each elbow points towards the TM38 of another Piezo1 subunit and in our model we found that Glu2133 on the elbows forms a salt bridge with the Arg2482 in a neighbouring subunit in agreement with previous suggestions^25^ (Fig. 3e). This salt bridge is retained in all our simulations between at least two Piezo1 chains with above 70% of occurrence, even after Piezo1 conformational rearrangement due to the applied negative pressure (Supplemental Table S2). This indicates that this interface is possibly maintained during the conformational transition we observed.

In agreement with this hypothesis, functional data^24^ demonstrated that the mutant G2133D increases the conductance with respect to the wildtype, which is probably due to the short chain of the aspartate that would pull backward (i.e towards the elbow) the TM38, thus reducing the pore constriction. Furthermore, this hypothesis rationalises the effect of the less efficient mutant E2133K that was shown to decrease Piezo1 conductance^24^. That is, the mutation to lysine opposite to the Arg2482 would push forward the TM38 due to repulsion, contributing in the increase of the channel constriction. In the E2133K mutant, the salt bridge is broken and so is the connection between the elbow and the TM38 which results in an impairment of the functional displacement connected to the blade expansion. In this condition, although less conducting, the channel is still functional^24^, which may be due to the aforementioned hydrophobic interactions between TM37 and TM38 in synergy with the proposed of lipid displacement which may act in concert. In conclusion, the predicted salt bridge between Glu2133 with Arg2482 in a neighbouring subunit would provide an interaction interface between the Piezo1 chains so that each blade extension would control the gating of a nearby Piezo1 chain (Fig. 4b). This may also provide a mechanism that ensures a prompt and mutual propagation of sensing between Piezo1 subunits.

In conclusion, this study reveals the molecular principles by which Piezo1 channel opens in response to increase membrane tension. The steps during Piezo1 opening are shown in Fig. 4b and are: 1) In a membrane without tension, Piezo1’s curved shape forces the bilayer to curve creating a characteristic indentation with a trilobed topology. This indentation is stable in native condition and the CED is hidden within it. In this inactive resting conformation, Piezo1 channel mouth is occupied by lipids. 2) Membrane tension on the bilayer results in the flattening and extension of its blades and tilting of its beams helices. 3) Piezo1 expansion and flattening allows the blades to “pull” helices 37 and 38, via hydrophobic interactions, to tilt and progressively move away from the channel pore opening Piezo1. 4) In the open conformation Piezo1 blades are fully extended and flat, CED is exposed, beam helices are almost parallel to the lipid bilayer. The elbow regions from neighbouring subunits, also pull the pore-lining helices 38. At the same time the increase in the area per lipid due to tension allows displacement of the lipids that occupy the channel mouth resulting in an open. Therefore, both conformational changes and the increase in the APL that allows movement of lipids outside the channel pore region are required for Piezo1 opening.

This work provides molecular understanding of the structural rearrangements within Piezo1 that lead to gating and highlight the critical role of membrane lipids in Piezo1 opening. Our study demonstrates how the fascinating fold and shape of Piezo channels allow this protein to sense mechanical force thus playing a central role in transduction of mechanical signals throughout the membrane. Our findings have important implications for understanding mechanosensing in general, which is critical in stem cell niche commitment, vascular biology and development and are instrumental for understanding Piezo1 channel-related disease such as lymphoedema, anaemia and malaria.

## METHODS

### Modeling the Piezo1 trimer

Structural data were obtained from the cryo-EM structure PDB: 6B3R^12^. Missing residues were added with MODELLER (v 9.19)^26^ and the loop refinement tool^27^ was used to remove a knot in one chain between residues 2066-2074. The best loop was selected out of 10 candidates according to the discrete optimized protein energy method^28^. The final Piezo1 model does not comprises the first 576 residues because they are not present in the template and residues 718-781, 1366-1492, 1579-1654, 1808-1951 which are exposed to the cytosol and predicted as unstructured. Therefore, each chain is composed by five non-overlapping fragments: residues 577-717, 782-1365, 1493-1578, 1655-1807 and 1952-2547. The Piezo1 model obtained was further energy minimised in vacuum with GROMACS 2016^29^ prior simulations.

### Coarse-grained simulations

Prior atomistic simulations the Piezo1 model obtained as described above was converted to a CG resolution and energy minimised. The coarse-grained molecular dynamics (CG-MD) simulation was essential to equilibrate the lipid bilayer around the protein and reconstitute the Piezo1 membrane indentation. The CG-MD simulations were performed using the Martini 2.2 force field^30^ and GROMACS 2016^29^. To model the protein secondary and tertiary structure an elastic network model with a cut-off distance of 7 Å was used. The elastic network restricts any major conformational change within the protein during the CG-MD simulations. For the equilibration simulation Piezo1 model was inserted in a complex asymmetric bilayer using the INSert membrANE tool^31^. Three independent system assembly steps and CG equilibrations were carried out, the first from both were used in this study. The composition of the model bilayer is the following: For the outer leaflet, 1-palmitoyl-2-oleyl-phosphtidylcholine (POPC) 55%, sphingomyelin (SM) 5%, 1-palmitoyl-2-oleyl-phosphtidylethanolamine (POPE) 20% and cholesterol 20%. For the inner leaflet, POPC 50%, POPE 20%, 1-palmitoyl-2-oleyl-phosphtidylserine (POPS) 5%, cholesterol 20% and phosphatidylinositol 4,5-bisphosphate (PIP_2_) 5%. The system was neutralised with a 150 mM concentration of NaCl. The model was further energy minimised and subsequently equilibrated for 500 ns with the protein particles restrained (1000 kJ · mol^-1^ · nm^-2^) to allow the membrane bilayer to equilibrate around the model. The equilibration was performed at 323 K, with protein, lipids and solvent separately coupled to an external bath using the v-rescale thermostat ^32^ (coupling constant of 1.0). The temperature of 323 K is above the transition temperatures of all lipid species in the systems, therefore avoiding the lipids to undergo phase transitions to the gel phase. Pressure was maintained at 1 bar (coupling constant of 1.0) with semi-isotropic conditions and compressibility of 3 × 10^−6^ using the Berendsen barostat ^33^. Lennard-Jones and Coulombic interactions were shifted to zero between 9 and 12 Å, and between 0 and 12 Å, respectively.

### Atomistic simulations

After the CG-MD equilibration the system was carefully checked. Two molecules of POPC were removed because the head-group was trapped between Piezo1 transmembrane bundles. Moreover, in order to restore membrane asymmetry, two molecules of PIP_2_ were removed because they flipped to the outer leaflet. The system obtained was minimised and further converted to atomistic resolution as described in ^34^. A concentration of 3.0 mM of Ca^2+^ (i.e. 50 ions) was added to the box and neutrality restored with counterions. The obtained system was energy minimised and subsequently equilibrated in four NPT ensemble runs of 20000 steps each with an increasing time-step from 0.2 fs to 2 fs and the protein particles restrained (1000 kJ · mol^-1^ · nm^-2^). A further equilibration step of 5 ns with a time-step of 1 fs and Cα atoms restrained (1000 kJ · mol^-1^ · nm^-2^) was performed in order to relax the Piezo1 model embedded in a highly curved bilayer (i.e. the membrane indentation). Stretch-induced conformational changes in the Piezo1 model were investigated by NPT ensemble unrestrained simulations of 50 ns were the bilayer plane (xy-plane) pressure was varied semiisotropically between -40, -30, -20, -5 and +1 bar, whereas the pressure in bilayer normal (z) direction kept at +1 bar. The +1 bar and the -30 bar systems were further extended to 100 ns. A similar approach revealed insightful for investigate structural transitions for the TREK-2 mechanosensitive channel16. All the atomistic systems were simulated using GROMACS 2016^29^ with CHARMM36 force field34 and a 2 fs time step. A Berendsen semi-isotropic pressure coupling^33^ at 1 bar was used during all the equilibration phases. The Parrinello-Rahman barostat^36^ was further used for the stretch-induced simulations. All simulations were performed at 323 K with protein, lipids and solvent coupled to an external bath using the v-rescale thermostat. Long-range electrostatics were managed using the particle-mesh Ewald method^37^. The LINCS algorithm was used to constrain bond lengths^38^.

### Analysis and computer graphics

To calculate the area of the box the tool gmx energy from GROMACS 2016^29^ was used to extract, for each frame, the x-side and the y-side of the simulated box. The area was subsequently calculated by multiplying these values as x-side × y-side. The root mean-square deviation was calculated on the protein Cα using the gmx rms tool from GROMACS 2016^29^. For each system the initial coordinates were considered as a reference structure. For the system at -40 bar, the error is the standard deviation. The A_proj_ was calculated as follow: The program visual molecular dynamics 1.9.3 (VMD)^39^ (http://www.ks.uiuc.edu/Research/vmd/) was used to calculate distances between the residue Ala641, located within the first bundle embedded in the bilayer, from each Piezo1 chain and for each trajectory. Values were then averaged by chain with the standard deviation as error. The average distance calculated was then used to calculate the radius *r* of the circumscribed circle using an in-house script dividing the average distance (i.e. the side of a regular triangle) by square root 3. The area of the circumference was then calculated as π × *r*^2^. In every step the standard deviation was calculated. The average partial density was calculated using the GROMACS 2016^29^ tool gmx density. The tilting angle relative to the bilayer normal and distance from the Piezo1 pore of the Piezo1 beams, pore-lining helices 38 and 37, were calculated using gmx bundle and gmx distance tools from GROMACS 2016^29^. For all calculations, the Piezo1 pore region comprised the same residues that were used for the calculation of the cavity (see below). The tool trj_cavity^40^ (https://sourceforge.net/projects/trjcavity/) was used for the calculation of the volume of the Piezo1 pore cavity as well as for the water molecules that overlap to it. The considered residues were 2450-2547, 2214-2224 and 2326-2334. The trajectories were fitted on the Cα of the considered residues before the calculation and options -dim 4 (i.e. the degree of how buried they were) and -spacing 1.4 (i.e. the size of the grid voxel in Å) were used. Salt bridges were calculated with VMD 1.9.3^39^ using a 6 Å cut-off and, when indicated (see Supplemental Table S2), without considering the first 30 ns of simulation. For the calculation of the APL the tool GridMAT-MD^41^, version 2.0 http://www.bevanlab.biochem.vt.edu/GridMAT-MD/) was used. The lateral pressure *P*_*L*_ was calculated as explained for the TREK-2 mechanosensitive channel^16^. Molecular graphics were done using VMD 1.9.3^39^. Data were plotted using Grace (http://plasma-gate.weiz-mann.ac.il/Grace/).

## ACKNOWLEDGEMENTS

ACK and DDV are funded by an Academy of Medical Sciences and Wellcome Trust Springboard Award. DJB is funded by a Wellcome Trust Investigator Award.

## AUTHOR CONTRIBUTION

D.DV. Built the model, performed simulations and analysed the data. The research was conceptualised by A.C.K. and D.J.B., A.C.K and D.J.B. acquired funding and supervised the project. D.DV. Wrote the first draft. All authors revised the first draft.

## ADDITIONAL INFORMATION

A supplemental figure (Supplemental Fig. S1) and two tables of additional data (Supplemental Table S1 and Supplemental Table S2) accompany this paper.

## DATA AVAILABILITY

Data will be available a doi at the University of Leeds

---

**Supplemental Figure S1.**
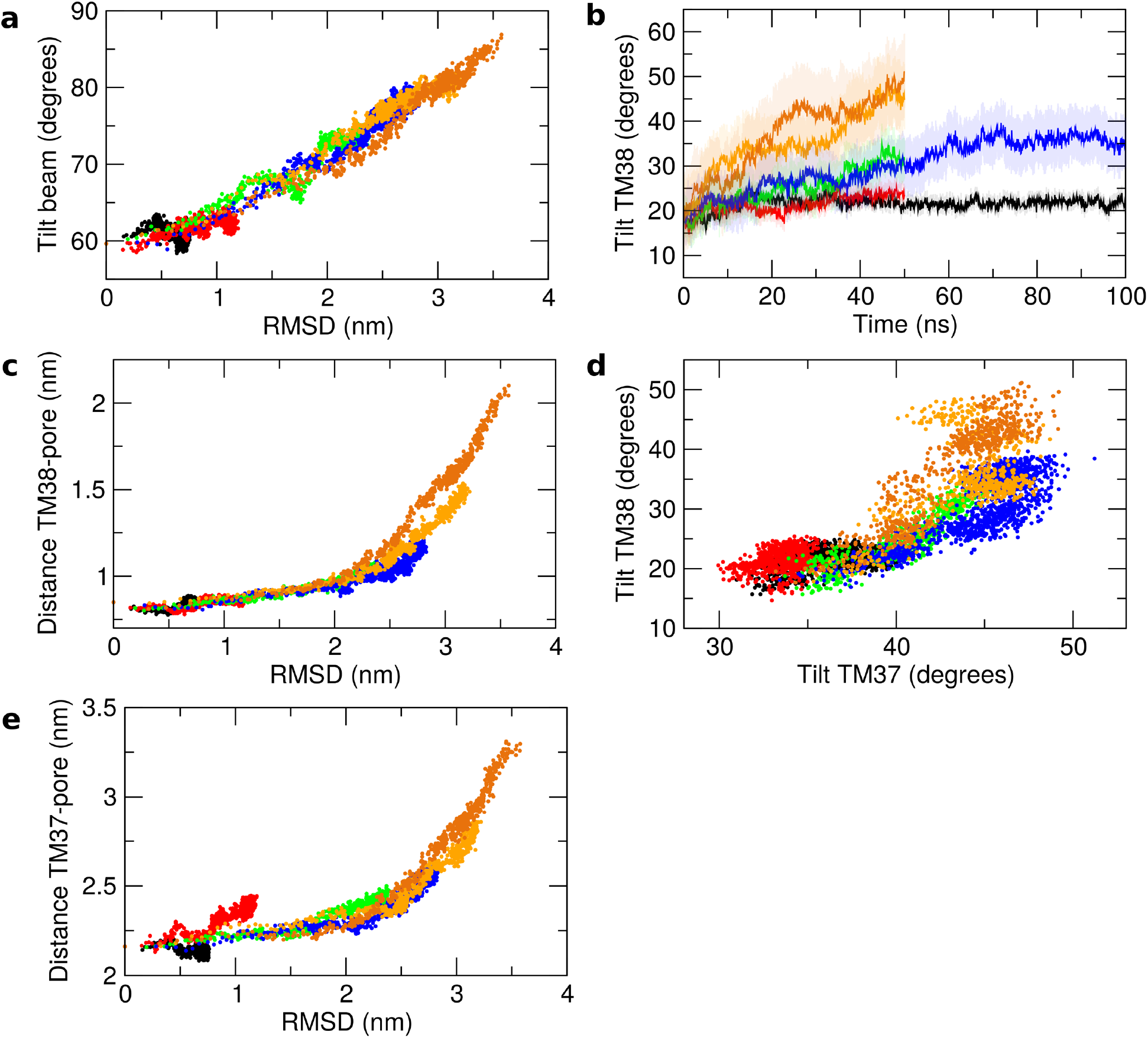
Correlation of Piezo1 pore-lining helices with changes in Piezo1 conformation. For each system, the applied negative pressure in each simulation is indicated with different colours: black, 1 bar; red, -5 bar; green, -20 bar; blue, -30 bar; orange, -40 bar. (**a**) Correlation between the root mean-square deviation (RMSD) of the Piezo1 blades and the tilt angle relative to the bilayer normal of the beam helix.(**b**) Tilt angle relative to the bilayer normal for the TM38 helix. (**c**) Correlation between the RMSD of the Piezo1 blades and the distance from the centre of mass of the TM38 helix and the Piezo1 pore. (**d**) Correlation between the tilt angle relative to the bilayer normal of the TM38 and TM37 helices. (**e**) Correlation between the RMSD of the Piezo1 blades and the distance from the respective centre of mass of the TM37 helix and the Piezo1 pore.

**Supplemental Table S1.**
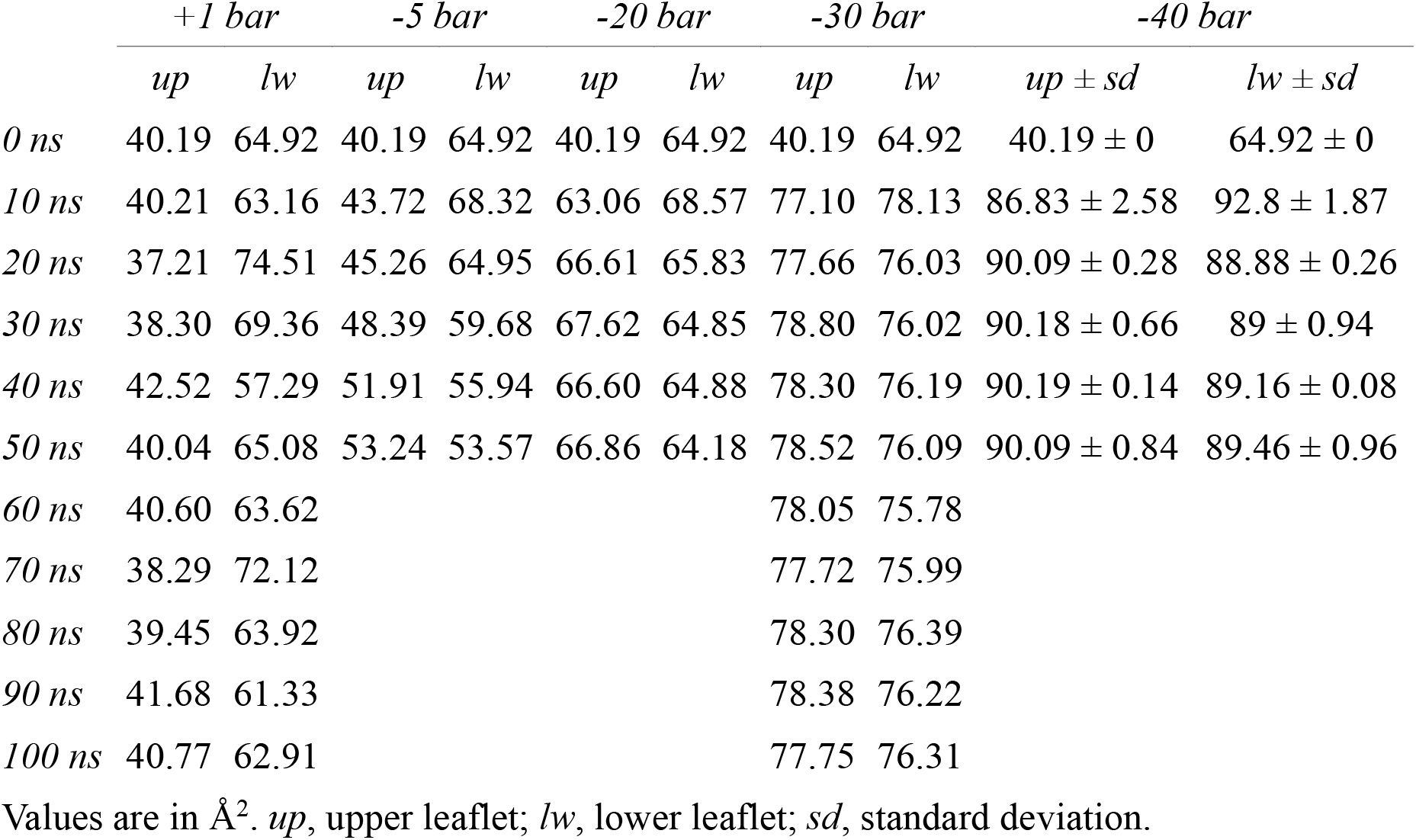
Area per lipid during simulations.

|  | +1 bar |  | -5 bar |  | -20 bar |  | -30 bar |  | -40 bar |  |
| --- | --- | --- | --- | --- | --- | --- | --- | --- | --- | --- |
|  | *up* | *lw* | *up* | *lw* | *up* | *lw* | *up* | *lw* | *up* ± *sd* | *lw* ± *sd* |
| 0 ns | 40.19 | 64.92 | 40.19 | 64.92 | 40.19 | 64.92 | 40.19 | 64.92 | 40.19 ± 0 | 64.92 ± 0 |
| 10 ns | 40.21 | 63.16 | 43.72 | 68.32 | 63.06 | 68.57 | 77.10 | 78.13 | 86.83 ± 2.58 | 92.8 ± 1.87 |
| 20 ns | 37.21 | 74.51 | 45.26 | 64.95 | 66.61 | 65.83 | 77.66 | 76.03 | 90.09 ± 0.28 | 88.88 ± 0.26 |
| 30 ns | 38.30 | 69.36 | 48.39 | 59.68 | 67.62 | 64.85 | 78.80 | 76.02 | 90.18 ± 0.66 | 89 ± 0.94 |
| 40 ns | 42.52 | 57.29 | 51.91 | 55.94 | 66.60 | 64.88 | 78.30 | 76.19 | 90.19 ± 0.14 | 89.16 ± 0.08 |
| 50 ns | 40.04 | 65.08 | 53.24 | 53.57 | 66.86 | 64.18 | 78.52 | 76.09 | 90.09 ± 0.84 | 89.46 ± 0.96 |
| 60 ns | 40.60 | 63.62 |  |  |  |  | 78.05 | 75.78 |  |  |
| 70 ns | 38.29 | 72.12 |  |  |  |  | 77.72 | 75.99 |  |  |
| 80 ns | 39.45 | 63.92 |  |  |  |  | 78.30 | 76.39 |  |  |
| 90 ns | 41.68 | 61.33 |  |  |  |  | 78.38 | 76.22 |  |  |
| 100 ns | 40.77 | 62.91 |  |  |  |  | 77.75 | 76.31 |  |  |

Values are in Å<sup>2</sup>. *up*, upper leaflet; *lw*, lower leaflet; *sd*, standard deviation.

**Supplemental Table S2.** Predicted salt bridge between the elbow and the pore-lining helix 38

|  | +1 bar | -5 bar | -20 bar | -30 bar | -40 bar<br>(rep1) | -40 bar<br>(rep 2) |
| --- | --- | --- | --- | --- | --- | --- |
| E2133.a - R2482.c | 100 | 100 | 100 | 100 | 100 | 100 |
| E2133.a - R2482.b | 98 | 98 | 100 | 96 | 70 (98) | 100 (100) |
| E2133.b - R2482.c | 100 | 100 | 100 | 7 (62) | 8 (89) | 7 (83) |

When occurrence was lower than 70%, numbers between parentheses indicate occurrence within the first 30 ns of simulations.

## REFERENCES

1. Wu, J., Lewis, A. H. & Grandl, J. Touch, Tension, and Transduction – The Function and Regulation of Piezo Ion Channels. Trends Biochem. Sci. 42, 57–71 (2017).

2. Beech, D. J. & Kalli, A. C. Force Sensing by Piezo Channels in Cardiovascular Health and Disease. Arterioscler. Thromb. Vasc. Biol. ATVBAHA119313348 (2019). doi: 10.1161/ATVBAHA.119.313348

3. Li, J. et al. Piezo1 integration of vascular architecture with physiological force. Nature 515, 279–282 (2014).

4. Fotiou, E. et al. Novel mutations in PIEZO1 cause an autosomal recessive generalized lymphatic dysplasia with non-immune hydrops fetalis. Nat. Commun. 6, 8085 (2015).

5. Andolfo, I. et al. Multiple clinical forms of dehydrated hereditary stomatocytosis arise from mutations in PIEZO1. Blood 121, 3925–3935 (2013).

6. Ma, S. et al. Common PIEZO1 Allele in African Populations Causes RBC Dehydration and Attenuates Plasmodium Infection. Cell 173, 443-455.e12 (2018).

7. Teng, J., Loukin, S., Anishkin, A. & Kung, C. The force-from-lipid (FFL) principle of mechanosensitivity, at large and in elements. Pflugers Arch. 467, 27–37 (2015).

8. Martinac, B., Adler, J. & Kung, C. Mechanosensitive ion channels of E. coli activated by amphipaths. Nature 348, 261–3 (1990).

9. Cox, C. D. et al. Removal of the mechanoprotective influence of the cytoskeleton reveals PIEZO1 is gated by bilayer tension. Nat. Commun. 7, 10366 (2016).

10. Poole, K., Moroni, M. & Lewin, G. R. Sensory mechanotransduction at membrane-matrix interfaces. Pflügers Arch. - Eur. J. Physiol. 467, 121–132 (2015).

11. Prager-Khoutorsky, M., Khoutorsky, A. & Bourque, C. W. Unique Interweaved Microtubule Scaffold Mediates Osmosensory Transduction via Physical Interaction with TRPV1. Neuron 83, 866–878 (2014).

12. Guo, Y. R. & MacKinnon, R. Structure-based membrane dome mechanism for Piezo mechanosensitivity. Elife 6, e33660 (2017).

13. Lin, Y.-C. et al. Force-induced conformational changes in PIEZO1. Nature 1–5 (2019). doi: 10.1038/s41586-019-1499-2

14. Wang, L. et al. Structure and mechanogating of the mammalian tactile channel PIEZO2. Nature 1–5 (2019). doi: 10.1038/s41586-019-1505-8

15. Murphy, E. J., Joseph, L., Stephens, R. & Horrocks, L. A. Phospholipid composition of cultured human endothelial cells. Lipids 27, 150–153 (1992).

16. Aryal, P. et al. Bilayer-Mediated Structural Transitions Control Mechanosensitivity of the TREK-2 K2P Channel. Structure 25, 708-718.e2 (2017).

17. Botello-Smith, W. M. et al. A mechanism for the activation of the mechanosensitive Piezo1 channel by the small molecule Yoda1. Nat. Commun. 10, 4503 (2019).

18. Wu, J. et al. Inactivation of Mechanically Activated Piezo1 Ion Channels Is Determined by the C-Terminal Extracellular Domain and the Inner Pore Helix. Cell Rep. 21, 2357–2366 (2017).

19. Coste, B. et al. Piezo1 and Piezo2 are essential components of distinct mechanically activated cation channels. Science 330, 55–60 (2010).

20. Zheng, W., Gracheva, E. O. & Bagriantsev, S. N. A hydrophobic gate in the inner pore helix is the major determinant of inactivation in mechanosensitive Piezo channels. Elife 8, e44003 (2019).

21. Borbiro, I., Badheka, D. & Rohacs, T. Activation of TRPV1 channels inhibits mechanosensitive piezo channel activity by depleting membrane phosphoinositides. Sci. Signal. 8, ra15 (2015).

22. Romero, L. O. et al. Dietary fatty acids fine-tune Piezo1 mechanical response. Nat. Commun. 10, 1200 (2019).

23. Pliotas, C. & Naismith, J. H. Spectator no more, the role of the membrane in regulating ion channel function. Curr. Opin. Struct. Biol. 45, 59–66 (2017).

24. Coste, B. et al. Piezo1 ion channel pore properties are dictated by C-terminal region. Nat. Commun. 6, 7223 (2015).

25. Kefauver, J. M. et al. Structure of the mechanically activated ion channel Piezo1. Nature 554, 481–486 (2017).

## METHODS ONLY REFERENCES

26. Šali, A. & Blundell, T. L. Comparative protein modelling by satisfaction of spatial restraints. J. Mol. Biol. 234, 779–815 (1993).

27. Fiser, A., Do, R. K. & Sali, A. Modeling of loops in protein structures. Protein Sci. 9, 1753–73 (2000).

28. Shen, M.-Y. & Sali, A. Statistical potential for assessment and prediction of protein structures. Protein Sci. 15, 2507–24 (2006).

29. Abraham, M. J. et al. GROMACS: High performance molecular simulations through multi-level parallelism from laptops to supercomputers. SoftwareX 1–2, 19–25 (2015).

30. De Jong, D. H. et al. Improved parameters for the martini coarse-grained protein force field. J. Chem. Theory Comput. 9, 687–697 (2013).

31. Wassenaar, T. A., Ingólfsson, H. I., Böckmann, R. A., Tieleman, D. P. & Marrink, S. J. Computational Lipidomics with *insane* : A Versatile Tool for Generating Custom Membranes for Molecular Simulations. J. Chem. Theory Comput. 11, 2144–2155 (2015).

32. Bussi, G., Donadio, D. & Parrinello, M. Canonical sampling through velocity rescaling. J. Chem. Phys. 126, 14101 (2007).

33. Berendsen, H. J. C., Postma, J. P. M., van Gunsteren, W. F., DiNola, A. & Haak, J. R. Molecular dynamics with coupling to an external bath. J. Chem. Phys 81, 3684–3690 (1984).

34. Wassenaar, T. A., Pluhackova, K., Böckmann, R. A., Marrink, S. J. & Tieleman, D. P. Going backward: A flexible geometric approach to reverse transformation from coarse grained to atomistic models. J. Chem. Theory Comput. 10, 676–690 (2014).

35. Lee, S. et al. CHARMM36 united atom chain model for lipids and surfactants. J. Phys. Chem. B 118, 547–556 (2014).

36. Parrinello, M. & Rahman, A. Polymorphic transitions in single crystals: A new molecular dynamics method. J. Appl. Phys. 52, 7182–7190 (1981).

37. Darden, T., York, D. & Pedersen, L. Particle mesh Ewald: An N·log(N) method for Ewald sums in large systems. J. Chem. Phys. 98, 10089–10092 (1993).

38. Hess, B., Bekker, H., Berendsen, H. J. C. & Fraaije, J. G. E. M. LINCS: A linear constraint solver for molecular simulations. J. Comput. Chem. 18, 1463–1472 (1997).

39. Humphrey, W., Dalke, A. & Schulten, K. VMD: visual molecular dynamics. J. Mol. Graph. 14, 33–8, 27–8 (1996).

40. Paramo, T., East, A., Garzón, D., Ulmschneider, M. B. & Bond, P. J. Efficient characterization of protein cavities within molecular simulation trajectories: trj_cavity. J. Chem. Theory Comput. 10, 2151–2164 (2014).

41. Allen, W. J., Lemkul, J. A. & Bevan, D. R. GridMAT-MD: A grid-based membrane analysis tool for use with molecular dynamics. J. Comput. Chem. 30, 1952–1958 (2009).

